# Release of extracellular DNA by *Pseudomonas* species as a major determinant for biofilm switching and an early indicator for cell population control

**DOI:** 10.1101/2021.02.11.430776

**Authors:** Fatemeh Bajoul Kakahi, Juan Andres Martinez, Fabian Moreno Avitia, Samuel Telek, Daniel C. Volke, Nicolas T. Wirth, Pablo I. Nikel, Frank Delvigne

## Abstract

The different steps involved in biofilm formation have been the subjects of intensive researches. However, the very early cell decision-making process related to the switch from planktonic to sessile state still remains uncharacterized. Based on the use of *Pseudomonas putida* KT2440 and derivatives with varying biofilm-forming capabilities, we observed a subpopulation of cells bound to extracellular DNA (eDNA) in the planktonic phase, as indicated by propidium iodide (PI) staining. Strikingly, the size of this eDNA-bound/PI-positive subpopulation correlated with the overall biofilm forming capability of the bacterial population. This finding challenges the conventional view of phenotypic switching and suggests that, in *Pseudomonas*, biofilm switching is determined collectively based on the quantity of eDNA released in the supernatant. The whole process can be followed based on automated flow cytometry, and the appearance of PI-positive cells was considered as an early-warning indicator for biofilm formation. For this purpose, automated glucose pulsing was used successfully to interfere with the proliferation of PI-positive cells, resulting in a reduction of biofilm formation. This study provides insights into the collective determinants of biofilm switching in *Pseudomonas* species and introduces a potential strategy for controlling biofilm formation.

## 1. Introduction

The transition from a planktonic to a biofilm state in a bacterial population can be influenced by a variety of factors and is a complex process that can vary depending on the microbial species considered and the environmental conditions^1–3^. It has been suggested that a subpopulation of cells within a planktonic culture may play a role in initiating biofilm formation by binding to extracellular DNA (eDNA)^4^. These cells can further serve as pioneers that start the biofilm formation process by adhering to surfaces and facilitating the recruitment of other cells to form either a mono- or a multi-species biofilm. In this context, it has been recently suggested that the process of cell aggregation in the liquid phase is a critical step in the biofilm lifecycle^5,6^. These aggregates offer several fitness advantages over individual planktonic cells, including increased antibiotic resistance and improved surface colonization^6–8^. Bacterial aggregation is influenced by various factors, such as quorum sensing, eDNA, ions, and cationic polymers^9–12^. Altogether, these findings indicate that bacteria display multicellular behaviors in the liquid phase that are distinct from those observed in surface-bound bacterial communities. Accordingly, cell auto- (i.e., aggregation between cells belonging to the same species) and co-aggregation (i.e., aggregation between cells belonging to different species) have then been considered as an important update to the biofilm cycle model^3^.

The involvement of extracellular DNA (eDNA) in the formation of bacterial aggregates has been observed in various bacterial species ^13–16^. eDNA plays multiple roles within the aggregate, serving as a structural component, an energy and nutrition source, and a gene pool for horizontal gene transfer in naturally competent bacteria ^17^. Additionally, eDNA has the ability to bind to bacterial flagella, leading to increased hydrophobicity of the cell surface. This, in turn, promotes bacterial aggregation and enhances the stability of the biofilm structure^18–20^. *Pseudomonas putida* KT2440 is a non-pathogenic soil bacterium endowed with the ability to adapt to a large variety of physicochemical and nutritional niches^21,22^, and able to form biofilms depending on the environmental conditions^23^. In our work, by using *P. putida* KT2440 and derivatives exhibiting either enhanced or reduced biofilm formation capabilities, we observed a fraction of cells bound to eDNA in the planktonic phase. This observation was based on propidium iodide (PI) staining. More importantly, it was determined that the size of the fraction of eDNA-bound/PI-positive cells is correlated to the global biofilm formation capability of the cell population. This result is important since it suggests that biofilm switching does not follow the classical phenotypic switching mechanism where individual cells within the population decide to activate or repress gene circuits according to environmental cues^24–26^. Instead, biofilm switching in *Pseudomonas* species seems to be determined collectively based on the number of eDNA molecules released in the supernatant. Massive release of eDNA increases the probability of binding to cells and the number of eDNA-bound cells, in turn, increases the probability of cell aggregation and biofilm formation. In this work, we will use these characteristics to characterize the dynamics of *P. putida* population upon continuous cultivation based on automated flow cytometry (FC)^27^. In a second, reactive FC strategies^28,29^ will be implemented for acting on the cell phenotypic switching mechanisms associated with biofilm formation in an attempt to either reduce or enhance biofilm formation.

## 2. Results

### Biofilm switching is determined early in the planktonic phase by a subpopulation of cells binding to eDNA

We first investigated biofilm switching based on standard cultivations in shake flasks. In order to establish a possible link between early eDNA excretion, PI staining and biofilm formation, we used different strains of *P. putida* exhibiting different biofilm forming capabilities (**Figure 1A**). For this purpose, we genetically manipulated the bacteria by either reducing or increasing their biofilm-forming ability. Specifically, we knocked out the *lapA* gene, which encodes the initial attachment protein, resulting in a strain that is unable to attach to abiotic surfaces. This Δ*lap*A derivative exhibits significantly reduced biofilm formation. To increase biofilm formation, on the other hand, we transformed *P. putida* with plasmid pS638::DGC-244, generating strain called DGC, (di-guanylate cyclase). This strain contains a more active di-guanylate cyclase that elevates the internal concentration of cyclic-di-GMP, a critical regulator of biofilm formation. High levels of c-di-GMP (c-di-GMP), support an adhesive lifestyle, while low levels lead to a planktonic lifestyle^30^. As expected, the *P. putida* DGC strain, with increased capacity for c-di-GMP production, exhibited the highest biofilm formation capability. On the contrary, the wild-type strain *P. putida* KT2440 exhibited low biofilm forming capability, which was further decreased upon the deletion of *lapA*. These three *P. putida* strains (KT2440 and derivatives) were then cultivated in shaken-flask cultures and monitored for PI staining (based on flow cytometry analyses) and eDNA release (**Figure 1B-D**). For all the strains, we observed a progressive accumulation of eDNA in the extracellular medium, correlated with the presence of PI-stained cells in the population. For the *P. putida* DGC strain, this effect was further increased and cell aggregates were also observed during the cultivation (**Figure 1E-G**). The presence of such aggregates is informative about a possible transition to biofilm lifestyle. Taken altogether, these data suggest that it could be possible to use PI as a molecular probe for tracking cells in planktonic phase in an early decision-making process for biofilm formation. The correlation between PI staining and cell eDNA binding was further confirmed based on the use of eDNAse (**Supplementary note 1**).

**Figure 1:**
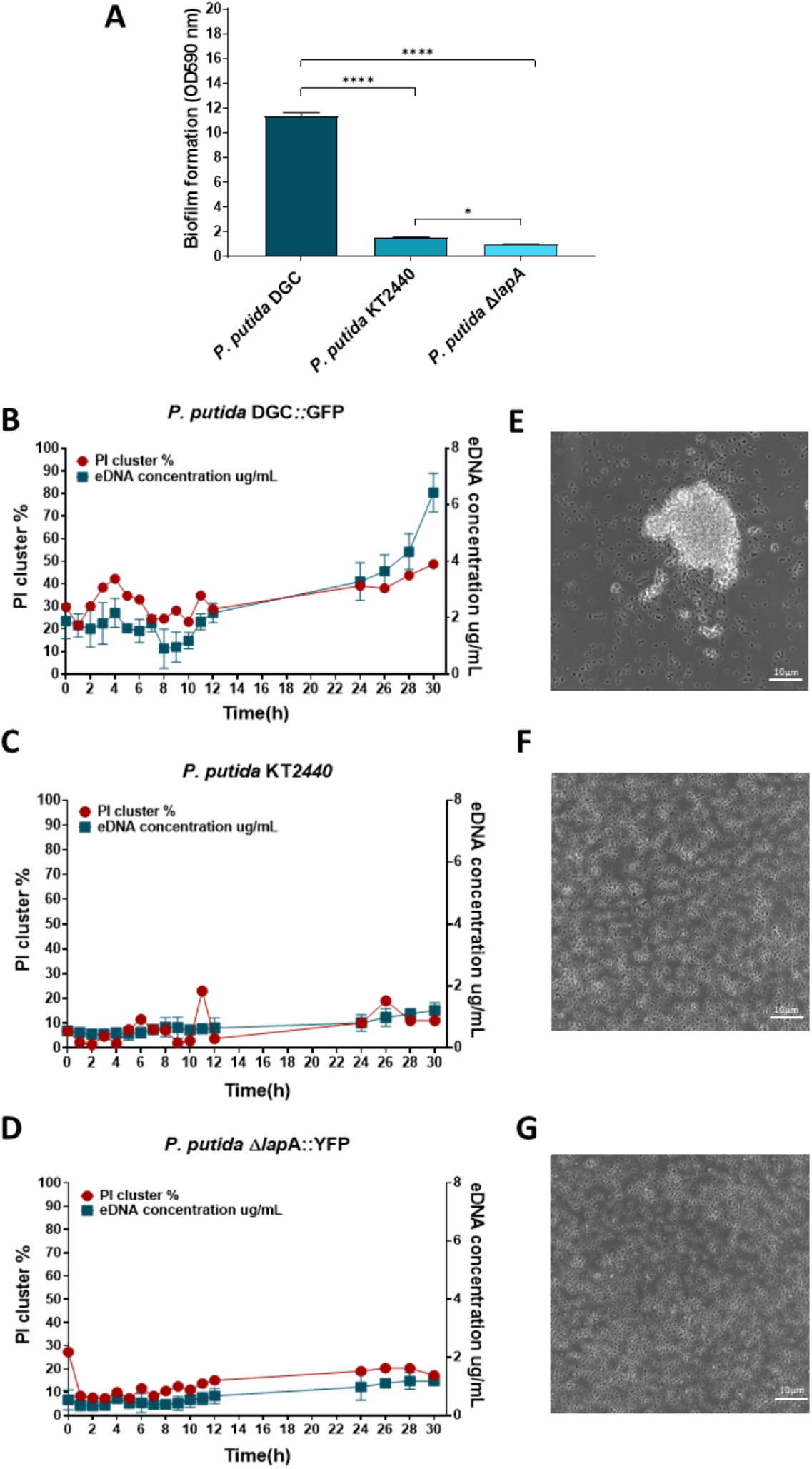
eDNA release increases the fraction of eDNA-bound cells in the planktonic phase. **A** Cristal violet assay for the biofilm forming capability of different derivatives of *P. putida* KT2440. Asterisks over brackets indicate a significant difference between samples (*p*-values: **** *p* < 0.0001, * *p* < 0.03). Results are presented as the mean and standard deviation for three biological replicates. **B-D** Time evolution of eDNA and PI positive cells (determined based on FC) during flask experiments (experiments have been done in triplicates). **E-G** Microscopy images (40x) of cells samples coming from flask experiments for *P. putida* DGC (**E**), *P. putida* KT2440 WT (**F**) and *P. putida* Δ*lapA* (**G**).

### eDNA binding is correlated with cell auto-aggregation

We used automated flow cytometry (FC) for mapping the population diversification dynamics in continuous culture. For this purpose, we use an in-house developed FC interface for sampling a chemostat at the time interval of 12 minutes^31^ (**Figure 2A**). In this way, we obtained a high-resolution temporal profile of the evolution of the population based on three parameters i.e., the red fluorescence associated with PI binding (FL3-A channel), as well as the size and morphology of the cells and cell aggregates (FSC-A and SSC-A parameters respectively). Indeed, we observed that the transition to the biofilm lifestyle in *Pseudomonas* was promoted by the formation of cellular aggregates in the planktonic phase (**Figure 1E** for *P. putida* KT2440 DGC and **Figure 2B&C** for *P. putida* KT2440 WT cultivated in chemostat). We then applied a specific data treatment procedure to the temporal profile obtained based on automated FC. In short, the time scatter plots related to the different signals i.e., PI (**Figure 2D**), FSC (**Figure 2E**), and SSC (**Figure 2F**), are divided into 50 fluorescence bins, each bin containing a specific number of cells. These fluorescence bins are used to compute the fluxes of cells into the phenotypic space by applying a gradient, leading to the quantification of the total fluxes of cells per time interval (F) for all signals (**Figure 2G-I**). The observation of the time evolution of the flux of cells through the diversification landscape reveals useful information i.e., that flux of diversification occurs stochastically during chemostat cultivation (**Figure 2J**). Interestingly, data analysis pointed out a strong direct relationship between the FC signals (**Figure 2K**), suggesting a close interplay between the PI signal and cell auto-aggregation, and subsequent biofilm formation.

**Figure 2:**
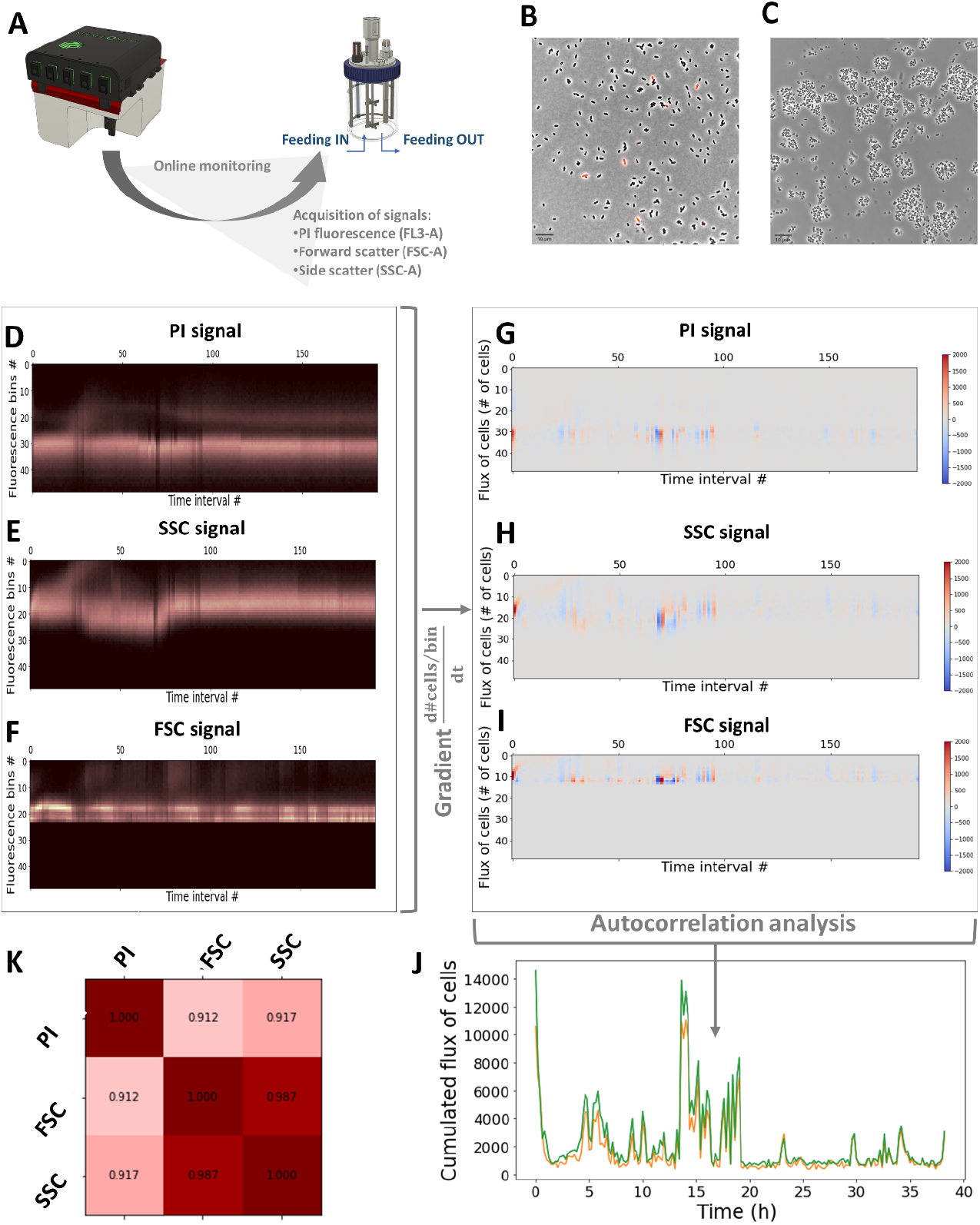
Automated FC analyses reveal a strong correlation between eDNA-bound, PI-positive cells and cell auto-aggregation. **A** Scheme of the automated FC set up connected to a continuous cultivation device (Dilution rate D = 0.1 h^-1^). **B-C** Microscopy images (40X) taken during continuous cultivation after 3h and 23h of cultivation respectively. **D-F** Binning of the time scatter profiles for the PI fluorescence, the FSC and SSC signals respectively (one time interval corresponds to 12 minutes). **G-I** computation of the gradient of cells from the binned data for the PI fluorescence, the FSC and SSC signals respectively (one time interval corresponds to 12 minutes). **J** Cumulated flux of cells extracted from the gradients of the three signals. **K** Correlation matrix with the Pearson coefficients indicating a strong direct relationship between the FC signals (p-values < 10^−78^ for all the correlations).

### Fast biofilm switching induces a fitness cost for planktonic growth

As stated in the previous section, fitness cost associated with phenotypic switching (or switching cost) is an essential factor affecting the diversification dynamics of the cell population. In a similar way, it can be expected that cells deciding to adopt a sessile lifestyle exhibit reduced fitness related to growth in the planktonic phase.

When we performed mono-culture with the three strains exhibiting different biofilm forming capabilities, we observed an inverse correlation between the fitness of the strain, as recorded based either on the global growth rate in (**Supplementary note 2**), or based on biomass yield in the planktonic (**Figure 3A**) and the biofilm phase (**Figure 3B**), and its biofilm forming capability. We further evaluated the switching cost by co-cultivating fast biofilm switcher (e.g., *P. putida* DGC) with strains exhibiting biofilm defect (i.e., *P. putida* Δ*lapA*). For ease of quantification, each strain was tagged with a different fluorescent protein. As expected, *P. putida* DGC::GFP was quickly outcompeted in the liquid phase in these conditions (**Figure 3C**) and was mainly found in the biofilm phase at the end of the cultivation (**Figure 3D**). The fact that biofilm switching induces a significant reduction in fitness for the planktonic phase will be exploited in the next sections for characterizing the diversification process of *P. putida* and designing biofilm mitigation strategies.

**Figure 3:**
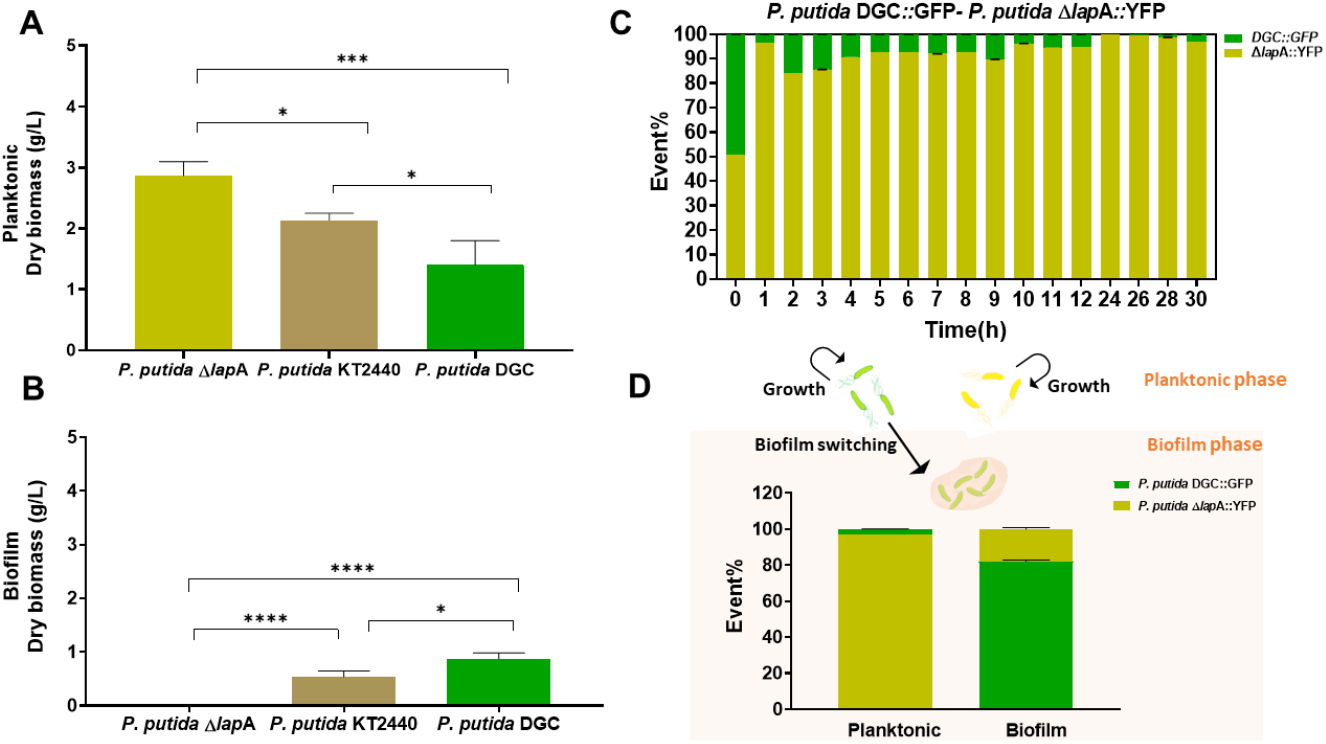
**A** Planktonic dry weight and **B** Biofilm dry weight comparison in mono- and co-cultures after 24h. The significance of the biofilm and planktonic dry weight data was evaluated (*p*-values: * *p* < 0.05, *** *p* < 0.0005, **** *p* < 0.0001).). **C** Dynamics of a co-culture *P. putida* DGC::GFP and *P. putida* Δ*lapA::*YFP cultivated in shake flask. Flow cytometry analysis of the percentage of each strain in the co-culture (n ≥ 3). The standard deviation has been computed from three biological replicates. **D** Flow cytometry analysis of the percentage of each strain in planktonic and biofilm phase after 24h of co-culture.

### Biofilm formation is associated with a bursty diversification dynamics

In microbial ecology, phenotypic switching is often associated with a fitness cost^25,32^. We recently demonstrated that this cost is the main driver for diversification dynamics of the whole population^27^. To illustrate this concept, we will take one of our previously characterized cellular system as an example. In the case of the phenotypic switching associated with sporulation in *Bacillus subtilis*, it is obvious that spore formation leads to a drastic reduction in growth for the cell deciding to switch. In this case, since the fitness cost associated with phenotypic switching is very high, a bursty diversification regime is observed. This regime is characterized by the appearance of spontaneous flux of cells (called bursts) phenotypically switching, these cells being progressively washed-out from the continuous cultivation device due to the loss of growth associated with the fitness cost. This specific diversification regime was previously reported during cultivation performed in a device called Segregostat^27^. In short, the Segregostat cultivation protocol relies on the use of reactive flow cytometry (**Figure 4A**) to detect the bursts of diversification (in a way similar to the determination of the fluxes of cells shown in **Figure 2J**). When a burst of diversification is detected, a pulse of glucose is added in order to interfere with the natural diversification process i.e., by giving a fitness advantage to the cells that are not differentiating. This concept is illustrated in the case of the sporulation in *Bacillus subtilis* (**Figure 4B**)^27^. In this case, a P_*spoIIE*_:GFP transcriptional reporter was used in order to detect the cells deciding to trigger sporulation. The bursts of cells can be visualized from the binned time scatter profile, as well as the concomitant fluxes of cells (**Figure 4C**). The observations made in the previous section point out that burstiness could also be associated with biofilm switching for *P. putida*. From a biological perspective, this would make sense since switching to the biofilm state, including cell auto-aggregation, results in a significant cost for the global growth of the population remaining in the liquid phase (**Figure 3**). However, a more complex diversification process has to be expected since population dynamics can also be impacted by cell auto-aggregation and cell release from biofilm (**Figure 4D**). The diversification profile of *P. putida* associated with biofilm switching was then determined based on Segregostat cultivation and PI staining, according to an automated protocol for PI staining previously established for different Gram-negative bacteria, including *P. putida*^31^. As expected, bursts of diversification were also observed in this case (**Figure 4E**). While the amplitude of these bursts is reduced compared to the ones observed upon standard chemostat cultivation (**Figure 2J**), they are more frequent and follow approximately the same periodicity as for the glucose pulses. In the next section, the detection of these bursts will be considered as an early-warning indicator of biofilm switching for *P. putida* KT2440. The Segregostat system will then be used to interfere with these bursts of diversification and to mitigate biofilm formation.

**Figure 4:**
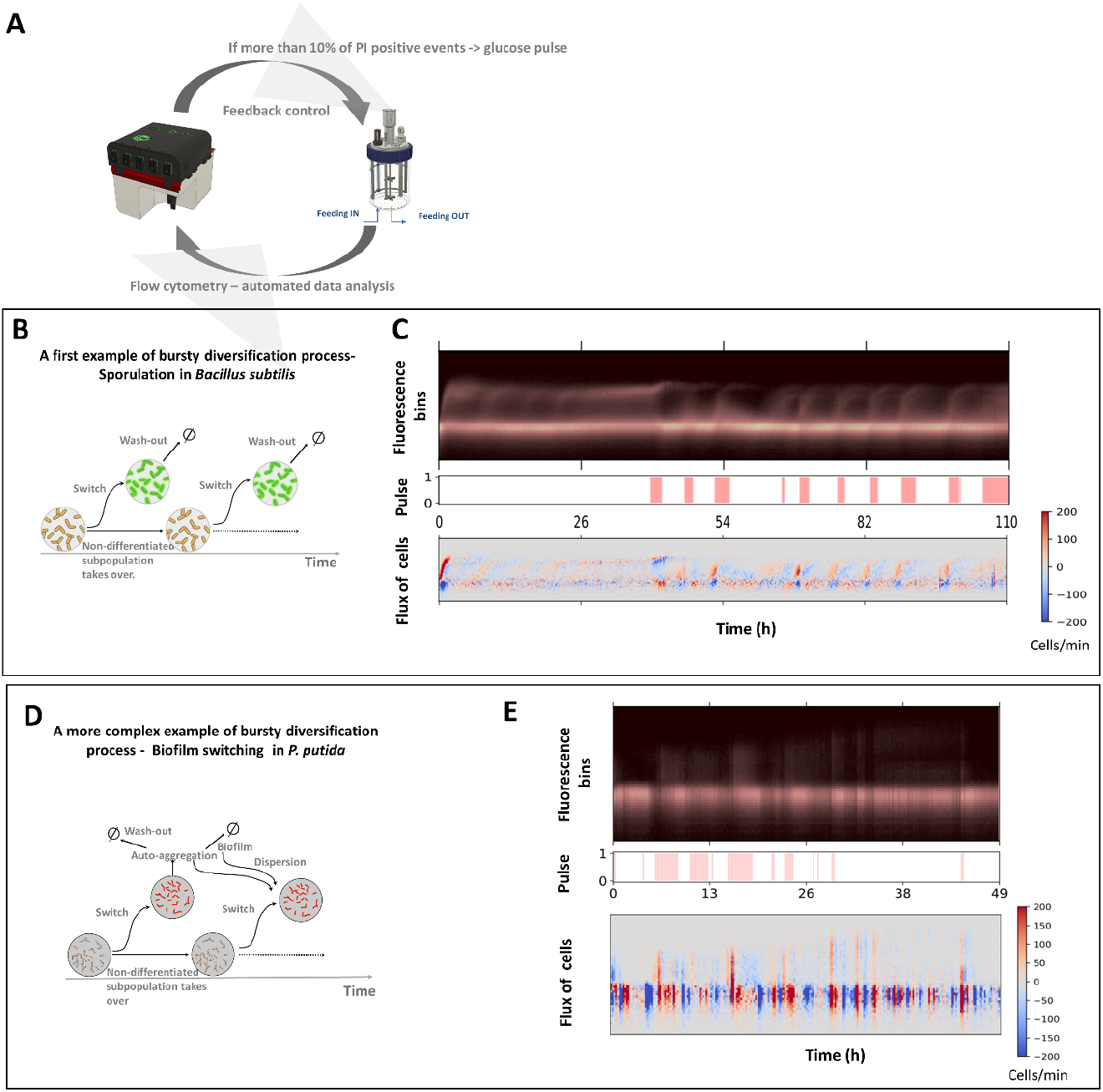
**A** Scheme of the Segregostat. Pulses of nutrients are added in the function of the ratio between PI negative and PI positive cells (here set at 10% of PI positive cells), as recorded by automated FC. **B** Example of bursty diversification process involved in the sporulation of *Bacillus subtilis*. **C** By applying environmental forcing based on Segregostat cultivation, the number of bursts is reduced, and the fluxes of cells involved in the process are increased, leading to a substantial but temporary reduction of the entropy for the population (adapted form Henrion et al 2023)^27^. **D** A more complex bursty diversification model proposed for *P. putida* KT2440. **E** Time evolution of fluorescence bins associated with PI staining. Flux of cells into the phenotypic space is computed by applying a gradient to the binned data. Time evolution of the total flux of cells into phenotypic space (The fluxes of cells in the phenotypic space F(t) have been computed from the binned fluorescence data).

### Control of bursts leads to reduced population diversification and biofilm formation

In the previous section, we observed that the diversification dynamics associated with biofilm switching was bursty. Our previous experiences with other bursty systems, like the sporulation in *Bacillus subtilis* or the T7-based expression system in *E. coli*, pointed out that these systems can be easily perturbed based on glucose pulsing since the differentiated fraction of cells exhibits a significantly reduced fitness by comparison with the non-differentiated one ^27^. This approach seems also to hold in the case of biofilm switching, given the fitness cost observed at this level (**Figure 3**). For this purpose, the diversification bursts were detected based on automated FC and glucose pulses were added according to a mode of cultivation called Segregostat. At this stage, two questions arise i.e., i. Does the Segregostat leads to a reduction of the global phenotypic heterogeneity of the population? and ii. Is it necessary to rely on automated FC for triggering glucose pulses, or can periodic pulsing be considered for this purpose?

In an attempt to reply to the first question, we previously used information theory for deriving a proxy that can be used for quantifying the degree of heterogeneity of a cell population i.e., the information entropy ^33^. The computation of the entropy profile H(t) can be done based on the same binning strategy as the one developed for computing F(t) (**Supplementary note 3**). The second question was challenged by comparing the Segregostat data to the one obtained in the chemostat and continuous culture where glucose pulses were periodically applied (**Figure 5A-C)**. Whereas the quantitative measurement of the total amount of biofilm inside the continuous bioreactor is challenging to determine, we can clearly see that, upon Segregostat cultivation, the global entropy profile H(t) is reduced by comparison to standard cultivation or when periodic pulsing is applied **(Figure 5D)**. Indeed, the computation of entropy reveals that, in the Segregostat, the microbial population is more homogenous and contains mainly PI negative cells, indicating that biofilm switching is less marked in this case.

**Figure 5:**
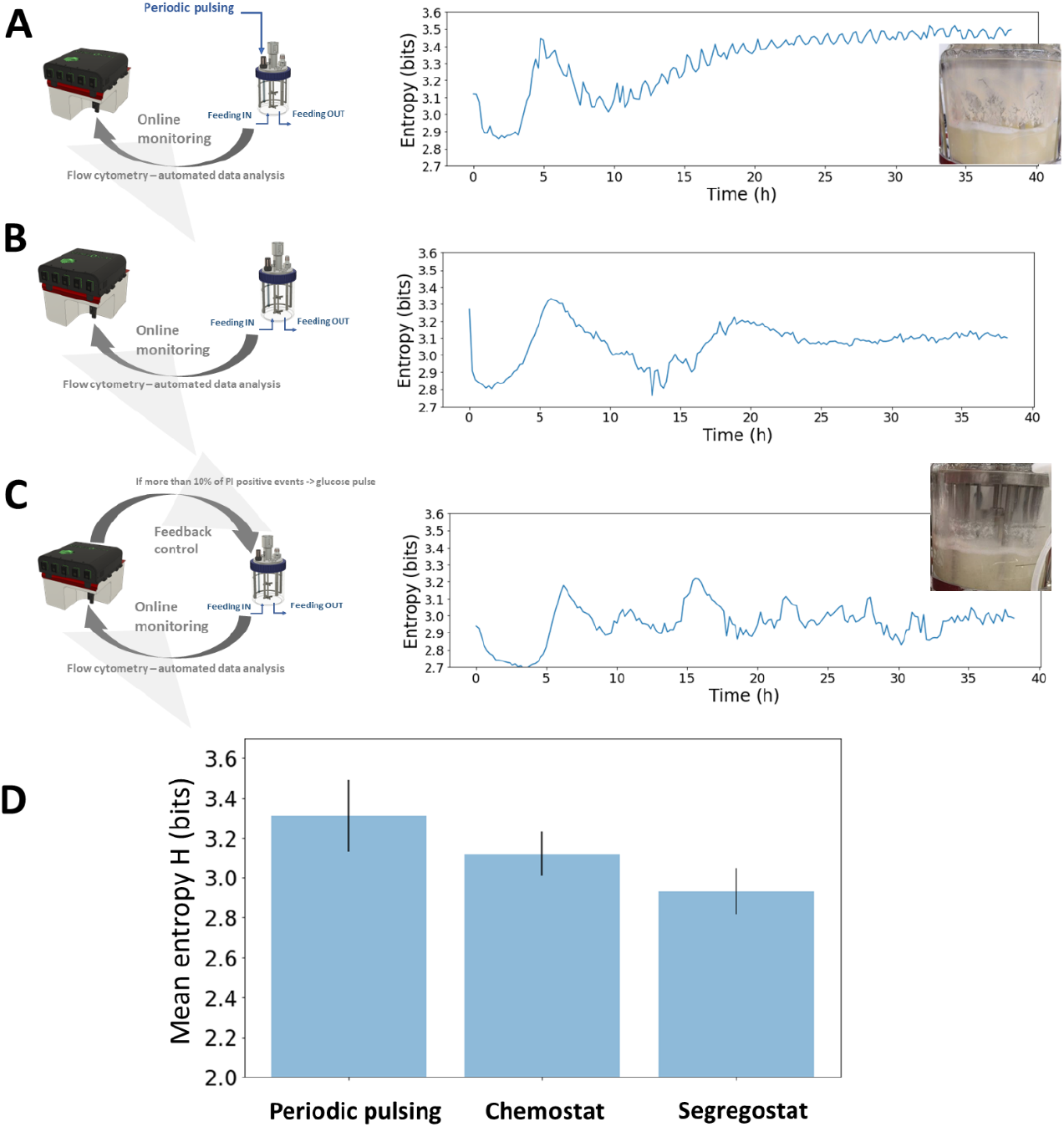
Evolution of the entropy H recorded in the function of the fitness cost associated to phenotypic switching in **A** Continuous cultivation at a dilution rate of 0.1 h^-1^ with periodic pulsing of glucose (0.5 g L^-1^ of glucose added each hour). **B** Chemostat cultivation at a dilution rate of 0.1 h^-1^ **C** Segregostat cultivation. Pictures of the biofilm formed on the well of the cultivation vessel are shown for the cultivations performed under periodic pulsing and Segregostat cultivations. **D** Comparative analysis of the mean level of information entropy H for the three modes of cultivation considered.

## 3. Discussion

Taken altogether, the data displayed in this work suggest that biofilm switching for *Pseudomonas* sp. is a collective decision taken by cells very early during planktonic growth, this decision being influenced by the amount of eDNA released to the extracellular medium. The transition from a planktonic to a biofilm state in bacterial populations is a complex process influenced by various factors and, accordingly, early cell decision-making process related to the switch to biofilm lifestyle has not been thoroughly investigated so far. However, this study suggests that a subpopulation of cells in planktonic cultures plays a pivotal role in initiating biofilm formation by binding to eDNA. Since these cells are bound to eDNA, a simple PI staining can be used for its detection. PI is commonly used in microscopy imaging to visualize extracellular DNA (eDNA) within biofilms or cell aggregates^4,34,35^. It has also been shown that the use of PI as a bacterial viability stain can yield misleading signals of cell death by binding to eDNA ^36^, a characteristic that can be exploited for detecting eDNA in biofilms^4^. We observed that the fraction of PI-positive/eDNA-bound cells is correlated with the overall biofilm formation capability of the population. This finding challenges the traditional view of phenotypic switching, where individual cells decide to activate or repress gene circuits based on environmental cues^26,32^. Instead, for *Pseudomonas* sp., biofilm switching appears to be determined on a collective basis, driven by the quantity of eDNA molecules released into the supernatant.

In Gram-negative bacteria like *Pseudomonas aeruginosa*, the release of extracellular DNA (eDNA) is mediated through various mechanisms. Quorum sensing, involving N-acyl-L-homoserine lactones (AHL) and the *Pseudomonas* quinolone signal (PQS), triggers eDNA release in planktonic cultures by inducing phage production^16^. Interestingly, recent findings have revealed that eDNA release in *P. aeruginosa* also occurs through oxidative stress caused by the generation of hydrogen peroxide (H2O2) mediated by pyocyanin production ^37^. Pyocyanin, a phenazine molecule that is regulated by quorum sensing ^38^, facilitates the binding of eDNA to *P. aeruginosa* cells. This association between pyocyanin and eDNA can impact the cell surface properties of *P. aeruginosa*, such as size, hydrophobicity, and surface energies, ultimately influencing cell-to-cell interactions and aggregation. Accordingly, a greater release of eDNA increases the probability of cell binding, which, in turn, promotes cell aggregation and biofilm formation^39^. Recent reports suggest that the presence of eDNA on bacterial cell surfaces promotes surface hydrophobicity and influences attractive acid-base interactions and therewith promotes initial bacterial adhesion and aggregation^40–42^. The dynamics of the process, as cells within the population switch from planktonic to biofilm state, has been characterized based on automated FC. FC offers a rapid and accurate method for quantifying specific cellular subpopulations within a biofilm^43^. Additionally, it has gained popularity as a means to investigate bacterial auto-aggregation^44,45^. PI-positive cells can indeed be easily determined based on FC and can also be used as an early-warning indicator for biofilm formation. Accordingly, we conducted a more precise characterization of biofilm switching based on automated flow cytometry (FC)^31^. Automated FC is indeed a valuable tool for monitoring the dynamics of cell population and is used here for tracking changes in the population as it transitions from planktonic to biofilm states^29^. The device can automate environmental transitions based on the phenotypic switching capability of the population^27,31^. In this work, we took benefit from Segregostat and the unique characteristics of eDNA to interfere with the biofilm switching mechanisms according to the detection of the burst of PI-positive cells. Furthermore, we demonstrated that using the environmental forcing implemented by Segregostat cultivation led to detection of the bursts of diversification and leads to a reduction in biofilm formation. Based on this experimental set-up, glucose pulses were administered in response to the detection of diversification bursts associated with PI-positive cells, giving a fitness advantage to non-differentiated cells in the liquid phase, thereby controlling phenotypic switching. The results showed that the Segregostat approach reduced the overall heterogeneity of the population, indicating a more homogenous microbial community with predominantly PI-negative cells, suggesting a decrease in biofilm switching.

In conclusion, this study provides valuable insights into the early determinants of biofilm switching in *Pseudomonas putida*, shedding light on the relationship between eDNA binding, PI staining, cell auto-aggregation, and biofilm formation. The findings offer new perspectives on microbial population dynamics and strategies for controlling biofilm formation. The application of Segregostat cultivation and the use of automated FC for monitoring and intervening in the diversification process open up possibilities for more targeted approaches to biofilm mitigation, with potential applications in various fields of microbiology and biotechnology^1,46^.

## 4. Material and methods

### Microbial strains and medium composition

The bacterial strains used in this study are listed in **Table 1**. All strains were maintained in 25% (v/v) glycerol at –80°C in working seed vials (2 mL). Prior to experiments, one colony of each bacterium was used to inoculate 10 mL of lysogeny broth (LB) medium (10 g L^-1^ NaCl, 5 g L^-1^ yeast extract, and 12 g L^-1^ tryptone) and grown for 6 h with shaking at 30°C. Precultures and cultures of all bacteria were done in modified M9 minimal medium (33.7 mM Na2HPO4, 22.0 mM KH2PO4, 8.55 mM 6 NaCl, 9.35 mM NH4Cl, 1 mM MgSO4, and 0.3 mM CaCl2), complemented with a trace element (13.4 mM EDTA, 3.1 mM FeCl3·6H2O, 0.62 mM ZnCl2, 76 μM CuCl2·2H2O, 42 μM CoCl2·2H2O, 162 μM H3BO3, and 8.1 μM MnCl2·4H2O), 1 μg L^-1^ biotin and 1 μg L^-1^ thiamin) and supplemented with glucose (5 g L^-1^) as the main carbon source (pH = 7.2). For strain DGC, the media was supplemented with gentamycin at a final concentration of 10 μg ml^-1^.

**Table 1.** Bacterial strains and plasmids were used in this study.

| Strain | Relevant characteristics<sup>a</sup> | Reference or source |
| --- | --- | --- |
| *E. coli* DH5α λpir | Cloning host; F<sup>-</sup> λ<sup>-</sup> *endA1 glnX44*(AS) *thiE1 recA1 relA1 spoT1 gyrA96*(Nal<sup>R</sup>) *rfbC1 deoR nupG* Φ80(*lacZ*Δ*M15*) Δ(*argF-lac*)*U169 hsdR17*(*rK*<sup>-</sup> *mK*<sup>+</sup>), λpir lysogen | 51 |
| *P. putida* KT2440 | Wild-type strain, derived from *P. putida* mt-2<sup>52</sup> cured of the TOL plasmid pWW0 | 53 |
| LapA | Derivate of *P. putida* KT2440 with a clean deletion of *lapA* (*PP_0168*) | This study |
| DGC | Derivate of *P. putida* KT2440 harboring the plasmid pS638::DGC-244 | This study |
| pSEVA638 | *oriV*(pBBR1); *XylS/Pm*→multiple cloning site (MCS); Gm<sup>R</sup> | 54 |
| pS638::DGC-244 | Derived from pSEVA638 with insertion of DGC-244 into the MCS | This study |
| pGNW2 | *oriV*(R6K); *P<sub>EM14g</sub>BCD*→*msfGFP*; Km<sup>R</sup> | 55 |
| pGNW2·Δ*lapA* | Derived from pGNW2 with homologous flanking region to *lapA* (*PP_0168*) | This study |
| pSEVA227M | Km<sup>R</sup>; *ori* (RK2), *Pem7* – *msf.GFP* | 56 |
| pSEVA227Y | Km<sup>R</sup>; *ori* (RK2), pBG<sub>17</sub>– YFP | 56 |

### Plasmid construction and genetic manipulations

Plasmid pS638::DGC was constructed by amplifying the hyperactive diguanylate cyclase mutant A0244 from *Caulobacter crescentus* ^47^, with the primer pair P1 and P2. The resulting amplicon and vector pSEVA638 were digested with *Bam*HI and *Sac*I (FD0054 and FD1133, Thermo Fisher Scientific). The fragments were purified (NucleoSpin Gel and PCR Clean-up Columns, Macherey Nagel) and ligated with T4 DNA ligase (EL0014, Thermo Fisher Scientific). *E. coli* DH5α was transformed with the ligation mixture and the cell suspension was plated on a gentamycin– selective plate. Subsequently, the constructed pS638::DGC (where *DGC*A0244 was placed under transcriptional control of the inducible XylS/*Pm* expression system) plasmid was isolated from a single colony and its correctness was confirmed by sequencing. The plasmids for the deletion of *lap*A were constructed according to ^48^. In short, ∼500 bp homology arms (HA) flanking the gene coding sequences were amplified from the chromosomal DNA of *P. putida* KT2440 using the primer pair P5/P6 (*lap*A_HA1) and assembled into the suicide vector pSNW2 employing the USER cloning method ^49^.The purified plasmids were introduced into stationary *P. putida* KT2440 cells by electroporation and selected on LB agar medium supplemented with kanamycin (50 μg mL^-1^). The corresponding pSNW2 derivative, now fully integrated into the bacterial chromosome, was resolved by transforming the cells with the auxiliary plasmid pQURE6 and selection on gentamicin (10 μg mL^-1^) and 3-methyl benzoate (1 mM) ^50^. Resolved strains (GFP-negative and kanamycin-sensitive) were tested for the desired genotype by colony PCR (OneTaq® 2X Master Mix with Standard Buffer, New England Biolabs). All the plasmids are listed in Table 2.

**Table 2.**
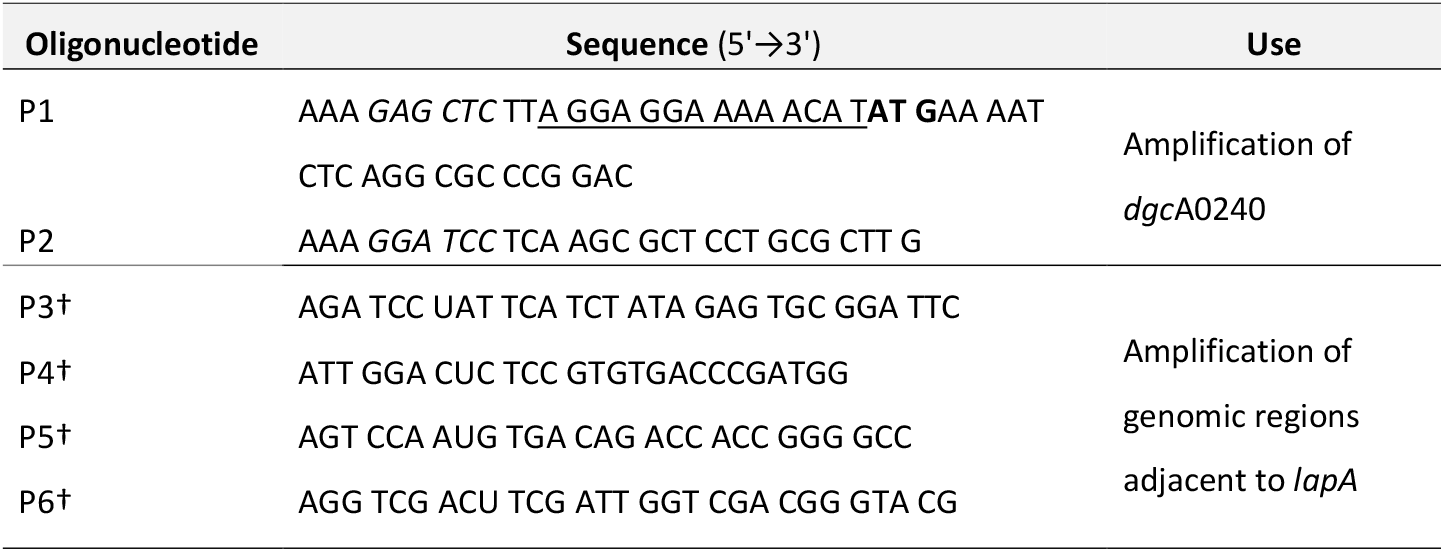
Oligonucleotides were used in this study. Oligonucleotides designed for USER cloning are indicated with a † symbol. Restriction sites are shown in italics, ribosomal binding site is underlined and start codon is shown in bold.

### Plasmid transformation

Plasmids pSEVA227M and pSEVA227Y, propagated in *E. coli* and *P. putida* KT2440 (kindly provided by Prof. Victor de Lorenzo, CNB-CSIC, Spain), were isolated with a plasmid extraction kit (NucleoSpin Plasmid EasyPure, Germany) according to the manufacturer’s protocol and stored at -20°C. Electrocompetent cells of *P. putida* Δ*lap*A were prepared according to a slightly modified procedure originally described by (Choi et al., 2006). Briefly, 1 ml of cells in the early stationary phase (OD600 = 1–1.5) from cultures grown in LB medium were harvested by centrifugation at 8000×g and washed twice with 1 ml of 300 mM sucrose at room temperature (RT). Cells were resuspended in 100 µl of 300 mM sucrose. In the case of *P. putida* DGC, electrocompetent cells were prepared by washing the biomass with 10% (v/v) glycerol. Briefly, 50 ml of a cell culture in the exponential phase (OD600 = 0.8) in LB medium were harvested by centrifugavon at 8000×g and washed twice with 50 ice-cold glycerol, centrifuge and resuspend cells in 0.8 ml ice-cold glycerol, keep on ice. Then 50 µl of electrocompetent cells were mixed with 1µl plasmid DNA (50 ng/ µl) in a 1 mm electroporation cuvette. High voltage electroporation was performed using a Gene Pulser Xcell (Bio-Rad Gene Pulser, US) at 25 µF, 200 Ω, and 1600 kV. After applying the pulse, 1 ml of SOC medium was added immediately, and the cells were transferred to a culture tube and incubated at 30 °C for 1 h. Cells were plated on LB agar plates supplemented with 50 μg/mL of kanamycin and incubated at 30 °C for 48–72 h.

### Quantification of biofilm growth in 96-well plates

Two-day biofilm formation of *Pseudomonas* strains was determined by crystal violet staining using a 96-well plate lid with pegs extending into each well (Nunc-TSP lid, Invitrogen™ Thermo Fisher Scientific). Briefly, precultures were grown overnight at 30°C to an OD600 nm of 1.0. The cell suspensions were then adjusted to an OD600 nm of 0.1 in M9 medium. A total of 160 μL cell suspension were added to each well. Fresh medium was used as a negative control. The plates were sealed with parafilm and incubated with shaking at 180 rpm at 30°C. The biofilm biomass was quantified with a crystal violet staining assay modified from previously reported CV assays ^57^. CV quantification was performed on the pegs of the Nunc-TSP lid culture system. Briefly, after 48 h of cultivation, the peg lids were taken out and washed three times using PBS. Subsequently, the peg lids were placed in plates with 180 μL of an aqueous 1% (w/v) CV solution. Then, the lids were washed with PBS three times after staining for 20 min. Subsequently, the peg lids with crystal violet stain were placed into a new microtiter plate with 200 μL of 33% (w/v) glacial acetic acid in each well for 15 min. The optical density at 590 nm of each sample was measured by a microplate reader (Tecan SPARK, Männedorf, Switzerland).

### Quantification of eDNA production in planktonic cultures

The bacterial strains, including *P*.*putida* KT2440, *P*.*putida* ΔlapA, and *P*.*putida* DGC, were cultured in modified M9 medium in 500mL Erlenmeyer flasks. These flasks were filled up to one-fifth of their nominal volume and placed on a rotary shaker at 170 r.p.m. at 30°C. The shake flasks were grown in parallel under identical conditions for a period of 30 hours, during which biomass growth and eDNA production were monitored. The cell density was estimated by measuring the OD600 nm (Genesys™ 10 UV-Vis, Thermo Scientific, USA). The eDNA in the supernatant was quantified after precipitation. To achieve this, bacterial cells were removed from 900 µL culture through centrifugation (4 min, 6800 g, Eppendorf™ 5424R). The supernatant (700 µL) was transferred to a sterile Eppendorf tube and mixed with 50 µL protein precipitation solution (Promega, USA) by inverting it ten times before centrifugation (10 min, 12100 g, Eppendorf™ 5424R). Then, 700 µL of the supernatant was mixed with 70 µL 2.5 M NaCl and 1400 µL 96% ethanol (62% final concentration) before being stored at -20 °C for at least 24 hours. The DNA was precipitated by centrifugation (25 min, 4 °C, 23 500 g, Eppendorf™ 5424R), after which it was washed once in 70% ice-cold ethanol and dried for less than 3 minutes at 37°C. The eDNA was quantified using the QuantiFluor dsDNA dye (QuantiFluor dsDNA System, Promega, Madison, WI, USA) according to the manufacturer’s protocol. In brief, eDNA in each sample was mixed with 200 μL of freshly prepared QuantiFluor dsDNA dye in TE buffer and incubated for 5 minutes before measuring the fluorescence intensity. The eDNA concentration was measured in 2 µL on a (NanoDrop™ 2000, Thermo Fisher Scientific, UK), using an excitation wavelength of 504 nm and an emission wavelength of 531 nm. For each run, a calibration curve was generated using Lambda DNA from Invitrogen™ Molecular Probes. To ensure the reproducibility of the experiment, the growth and eDNA concentration were determined in three independent samples for each time point. The experiment was also repeated with triplicates for all strains to verify the reproducibility between biological replicates.

### Biomass quantification

The dry matter of the liquid phase and biofilm phase were determined separately by the Moisture analyzer (HE53 Halogen Moisture Analyzer, Switzerland) according to manufacturer protocol. Briefly, the equipment was first warmed up for 30 minutes, and the standby temperature was set to 60°C. Then, the weighing aluminum pan containing a membrane filter disc (0.2µm, 47mm, Fisherbrand™) was tared automatically on the balance after being dried at 105°C in the moisture analyzer until reaching a stable weight lasting about 1 minute. The drying temperature was set to 105°C. Next, 10 mL of well-mixed planktonic and biofilm samples (After emptying the flask from planktonic culture, adherent cells were harvested by scraping from the wall of flasks with a cell scraper and resuspended in PBS and well-mixed) were evenly added to the membrane filter and positioned in a vacuum filtration apparatus. The liquid component of the sample was removed substantially by applying a vacuum from a small compressor for 2-5 minutes, leaving the broth solids. The filter and the residual solids were washed with 10 mL of deionized water, and the vacuum was reapplied to remove excess liquid. The filter and solids were replaced on an aluminum pan, and the drying program was set to end when the weight change was less than 0.1 mg min^-1^. The loss of weight upon drying was then used to calculate biomass as grams of dry weight per liter.

### Off-line flow cytometry analyses

FCM analysis was carried out with an Attune NxT Acoustic Focusing Cytometer (Thermo Fisher Scientific, United States) containing a violet laser 405 nm (50 mW), a blue laser 488 nm (50 mW), and a red laser 638 nm (100 mW). Instrument calibration was performed with Attune performance tracking beads (2.4 and 3.2 μm) (Thermo Fisher, United States). Side scatter (SSC) and Forward scatter (FSC) and BL3 (695/40) for PI (P4170, Sigma-Aldrich) were determined with FC. The software settings were as follows: Fluidics, medium; Threshold, 2000 on SSC-H; Run with limits, 40,000 events at a flow rate of 25 μl/min. Cells were diluted to an appropriate density OD600 (0.001-0.003 ≈700-1500 event/ μl) with filtered 1x PBS. FSC and SSC voltage and threshold were set based on wild-type bacteria. All bacterial samples were stained right before FC analysis by adding 1 μL of the PI stock solution (a stock solution was prepared at 1 mg mL^-1^ in sterile Milli-Q water and used at a final concentration of 1.5 μM in sterilized PBS) to 1 mL of a cell suspension in PBS (1x10^7^ cells mL^-1^). The stained samples were incubated for 10 min in the dark at room temperature and analyzed by FC (live-dead gating was done based on heat-killed bacteria at 80°C for 1 h).

### Microscopy imaging

Samples from *P. putida* KT2440, *P. putida* Δ*lap*A, and *P. putida* DGC culture were taken at 0h, 6h, and 24h during the batch phase and subjected to microscopy analysis. Microscopy images were acquired using a Nikon Eclipse Ti2-E inverted automated epifluorescence microscope (Nikon Eclipse Ti2-E, Nikon France, France) equipped with a DS-Qi2 camera (Nikon camera DSQi2, Nikon, France), a 100× oil objective (CFI P-Apo DM Lambda 100× Oil (Ph3), Nikon, France).

### Continuous cultivation with automated FC

In this study, *P. putida* KT2440 was grown in a stirred bioreactor (Biostat B-Twin, Sartorius) with a total volume of 2 L and a working volume of 1 L. The batch phase was initiated by diluting overnight cultures into a minimal medium to achieve an initial OD600 of 0.3. The culture was maintained at pH 7.2 and 30°C, with stirring set at 800 rpm and an aeration flow rate of 1 L min^-1^ (1 vvm). The depletion of oxygen marked the end of the batch phase and the start of continuous cultivation mode. For chemostat cultivations, the modified M9 medium containing 5 g L^-1^ glucose was continuously fed at a dilution rate of 0.1 h^-1^. In contrast, for periodic pulsing system cultivations, a modified M9 medium without a carbon source was continuously fed at the same dilution rate, and glucose feed pulses (0.5 g L^-1^ per pulse) were introduced hourly. The Segregostat platform, previously described^31,58^was used as a reactive FC strategy, in which glucose was pulsed based on a predefined set point. A feedback control loop, including a custom MATLAB script based on FC data, activated a pump to pulse an actuator (glucose) accordingly. The regulation was triggered once the fluorescence threshold was exceeded by more than 10% of the PI positive cells (detected based on the FL3-A channel). Throughout the experiments, samples were collected every 12 minutes from the bioreactor and automatically diluted and stained with PI and analyzed by flow cytometry (BD Accuri C6, BD Biosciences) based on a FSC-H threshold of 20,000.

Automated FC data were then processed according to custom Python codes (**Supplementary note 3**).

## Acknowledgement

JAM is supported by a post-doctoral grant provided by the Service Public de Wallonie (SPW) and the H2020 program of the European commission (Era-Cobiotech project Contibio). The financial support from The Novo Nordisk Foundation (NNF10CC1016517 and NNF18CC0033664) to P.I.N. is gratefully acknowledged.

## Notes

### Competing Interest Statement

The authors have declared no competing interest.

### Summary of Updates

The relationship between the eDNA-bound, PI-positive, subpopulation and biofilm switching has been more precisely established. Control of biofilm based on automated flow cytometry detection of the PI-positive cell population has been established and more thoroughly characterized.

