## Supplementary material for "Release of extracellular DNA by *Pseudomonas* species as a major determinant for biofilm switching and an early indicator for cell population control"

**Supplementary information**

### Supplementary note 1: DNAse treatment and confocal laser scanning microscopy (CLSM)

Planktonic cells were stained with PI and SYTO9, and it was observed that the two stains generally co-localized. Interestingly, within individual sections, green fluorescing cores were observed beneath the red-stained shells, indicating that PI staining was not indicative of membrane integrity, but rather the presence of eDNA outside the intact membrane. The results clearly demonstrated that PI signals were much more intense in the non-treated sample compared to the sample treated with DNase (**Supplementary Figure 1**).
Biofilm and planktonic cell densities were adjusted to ($1\times10^{7}$ cells $mL^{-1}$), and samples were resuspended in 500 µL of 1× DNase I buffer (10 mM Tris·HCl, pH = 7.5, 2.5 mM MgCl2, and 0.1 mM CaCl2) with or without DNase I (final concentration 160 U $mL^{-1}$, Roche, reference number 04716728001) and were incubated at 37°C 10 for 3 h. After incubation, samples were pelleted by centrifugation at 8500 rpm for 10 min, resuspended in PBS, stained by PI, and analyzed by FCM.
All samples were analyzed by CLSM with a LSM880 Airyscan super-resolution system (Carl Zeiss, Oberkochen, Germany). The images were taken based on a Plan- Apochromat 63×/1.4 Oil objective. We used an excitation wavelength of 488 nm and emission at 500-550 nm for green fluorescence and an excitation wavelength of 561 nm and emission at 580-18 615 nm for red fluorescence. Images were acquired continuously at a pixel resolution of 0.04 µm (regular Airyscan mode) in XY and 1-µm interval in Z step-size using the piezo drive. Before CLSM analysis, staining was performed with one or more fluorescent dyes i.e., either PI alone or a combination of SYTO9 and PI. An aliquot of 10-15 µL of 1:1 stain mixture solution was added to a sample of planktonic and biofilm and incubated under the exclusion of light 10 min.

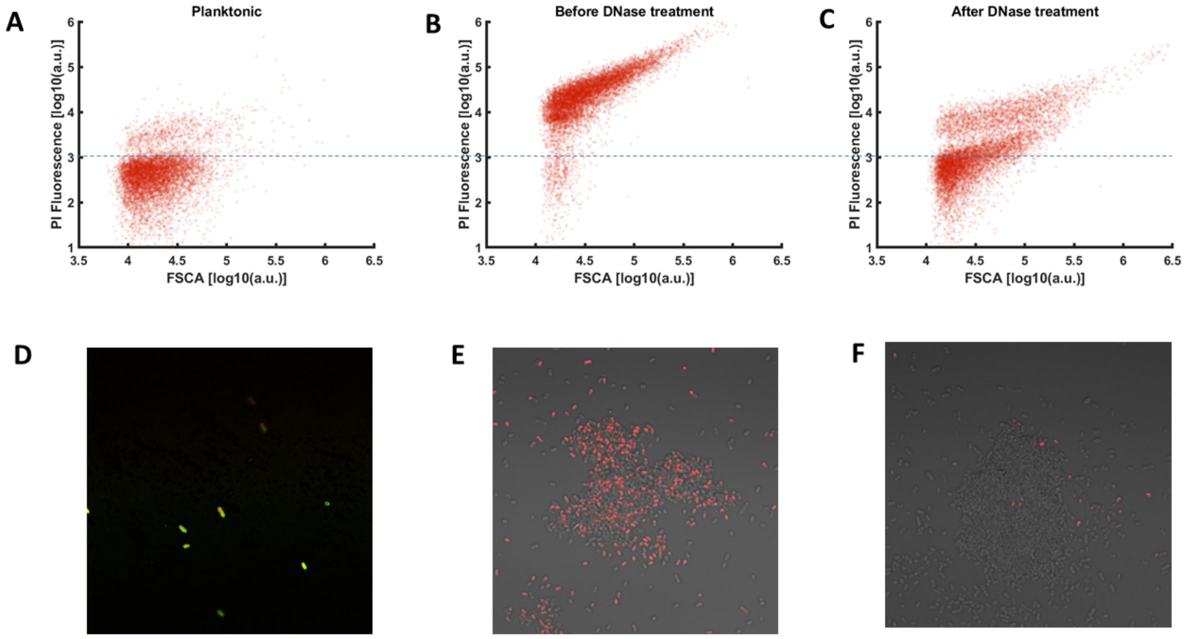


**Supplementary Figure 1**: **Flow cytometry analysis illustrating the comparison of PI positive percentage among planktonic and biofilm sample and effects of DNase treatment on PI positive fraction of biofilm sample**. **A** Planktonic sample co-stained with PI and SYTO 9. **B** Sample showing high PI uptake, before being treated by the DNase enzyme **C** Sample After treatment with DNase enzyme, the PI-positive fraction decreases significantly. Confocal laser scanning microscopy (CSLM)

images of *P. putida* DGC (Planktonic and biofilm) samples **D** Planktonic cells stained based on PI and SYTO 9. **E** Biofilm cells stained with PI before being treated with DNase enzyme. **F** Biofilm cells stained with PI after being treated with DNase enzyme.

#### Supplementary note 2: cultivation dynamics in shake flasks

The three strains of *P. putida* were cultivated in shake flask for determining their fitness in liquid culture and the fitness cost related to biofilm switching (**Supplementary Figure 2**). For this purpose, the time evolution of biomass and carbon source (glucose) were determined. Gucose concentrations were analyzed by high-performance liquid chromatography (Waters Acquity UPLC® H-Class System) using an ion exchange Aminex HPX-87H column (7.8 × 300 mm, Bio-Rad Laboratories N.V.). The analysis was carried out with an isocratic flow rate of 0.6 mL $min^{-1}$ for 25 min at 50°C. The mobile phase was composed of an aqueous solution of 5 mM H2SO4. Elution profiles were monitored through a Waters Acquity® Refractive Index Detector (RID) (Waters, Zellik, Belgium). Glucose standard solutions (Sigma-Aldrich, Overijse, Belgium) were used to determine the retention times and construct calibration curves.

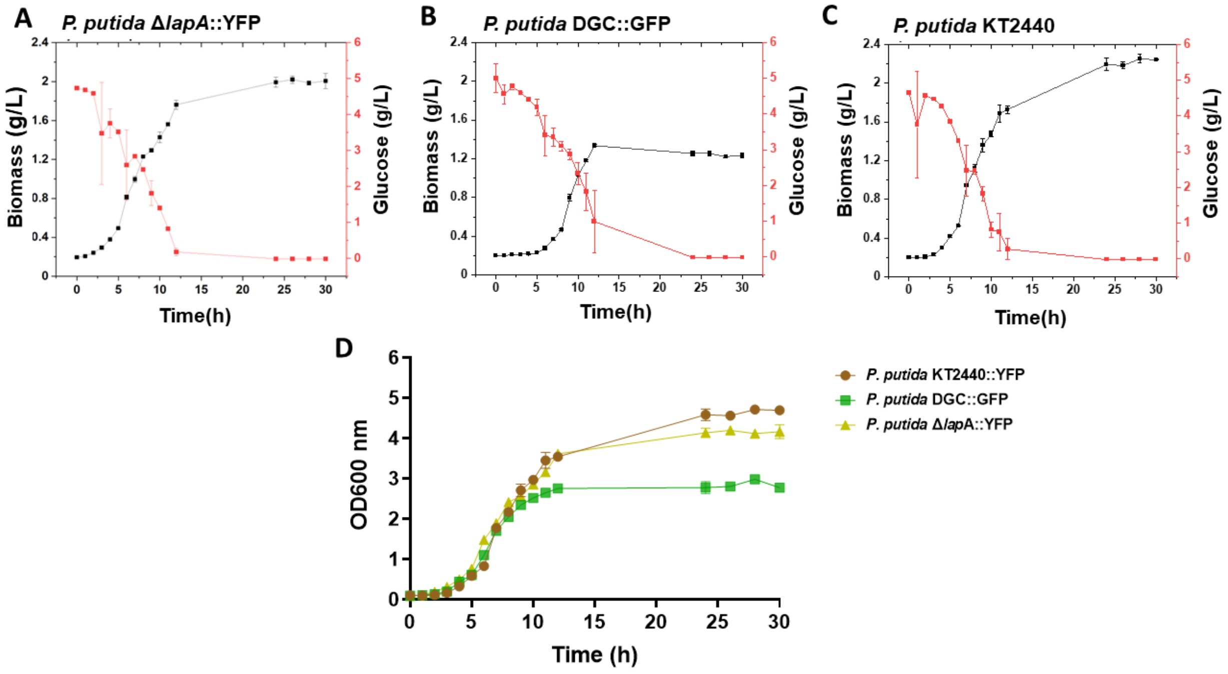


**Supplementary Figure 2: Evolution of biomass and glucose concentration during shake flask cultivation. A** *P. putida* Δ*lap*A::YFP , **B** *P. putida* DGC::GFP , **C** *P. putida* KT2440. **D** Comparison of growth curves of *P. putida* ΔlapA::YFP , *P. putida* DGC::GFP , *P. putida* KT2440 during 30h of batch cultivation, n ≥ 3 determined by spectrophotometric measurements (OD600). Each curve resembles a biological replication. The error bars represent the standard deviation (SD) based on three biological replicates.

**Supplementary note 3:** Processing the data from automated FC and computing of the flux of cells (F) and the entropy (H) of the cell population

Microfluidics are typically used for characterizing population dynamics with a single cell resolution[12]. However, microfluidics cultivation experiments, as well as the associated data processing, are quite time consuming. At this level, FC can be considered as an alternative method exhibiting a higher experimental throughput[3], but at the expense of losing the information about specific trajectories of the individual cells[4]. This drawback can be compensated by considering automated FC protocols where more regular snapshots of the population can be acquired. We used automated FC in the context of this work for tracking PI-positive, eDNA-bound, cells. Each 12 minutes, a sample was automatically taken out of the bioreactor, diluted, stained with PI and analyzed by FC. Individual snapshots can then be assembled into a time scatter plot (**Supplementary Figure 3A**).

Additional data treatment steps can also be applied for extracting information about the degree of heterogeneity of the cell population, as well as the fluxes of cells through the phenotypic space. This study employs a proxy derived from information theory i.e., information entropy H, to characterize how cell populations dispersion evolves with time[5]. Information theory is being more and more applied to study signal processing by cellular systems[678910]. The fundamental basis of information theory relies on the quantification of information entropy (H), serving as a gauge of uncertainty regarding the cell population's response (output) concerning environmental stimuli (input)[11]. However, the entropy profile H(t) will be used for characterizing the dynamics of population dispersion over time.
To quantify entropy, we utilize the following equation based on the PI fluorescence distribution acquired through automated flow cytometry (FC):

$$\textbf{Eqn 1}: \mathrm{H} = -\sum_{i=1}^{m} \mathrm{p}(x_i).\log_2 \mathrm{p}(x_i)$$

Here, m represents the number of observed states (PI classes in our case), and p denotes the probability of observing a cell displaying a specific state i.e., a given PI fluorescence intensity in our case. The probabilities for various fluorescence classes are easily determined based on automated FC. For this purpose, the PI distribution for a given interval of time is divided into 50 different bins (corresponding to m = 50 in **Equation 1**) and the entropy is determined based on the specific number of cells inside each bin by applying **Equation 1** (**Supplementary Figure 3B**). The number of bins has been previously shown to be optimal for determining the entropy based on FC data[12].

The binned data can also be used for determining the flux of cells (F) through the phenotypic space. For this purpose, a gradient is applied over 2 consecutive binned datasets (**Supplementary Figure 3C**). Based on the computation, the F(t) profile can be determined.

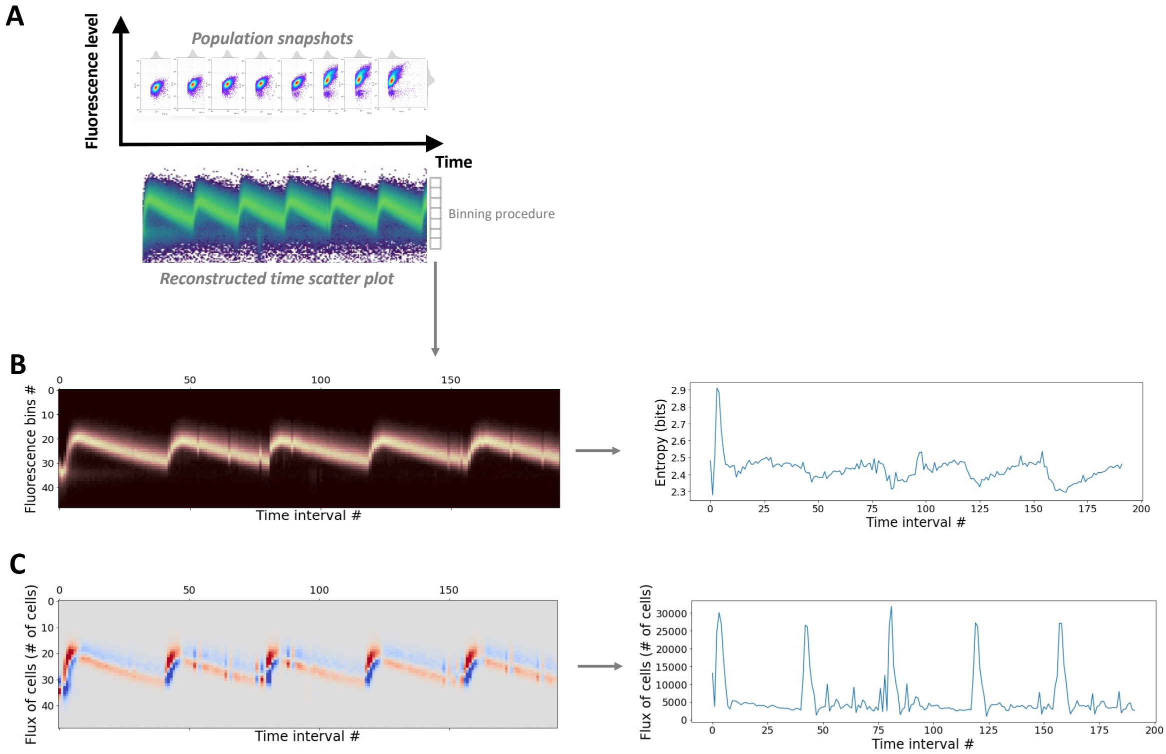


**Supplementary Figure 3: Data processing steps related to automated FC. A** Automated FC allows for the acquisition of snapshots of the population at regular time intervals (12 minutes). These snapshots can be assembled into a time scatter profile. **B** Binning is applied to each snapshot of the time scatter profile for computing the evolution of information entropy H(t). **C** Binned data can be further processed for determining the flux of cells based on the enrichment or depletion of each bin.
